# Condensin extrudes DNA loops in steps up to hundreds of base pairs that are generated by ATP binding events

**DOI:** 10.1101/2020.11.04.368506

**Authors:** Je-Kyung Ryu, Sang-Hyun Rah, Richard Janissen, Jacob W. J. Kerssemakers, Andrea Bonato, Davide Michieletto, Cees Dekker

## Abstract

The condensin SMC protein complex organizes chromosomal structure by extruding loops of DNA. Its ATP-dependent motor mechanism remains unclear but likely involves steps associated with large conformational changes within the ~50 nm protein complex. Here, using high-resolution magnetic tweezers, we resolve single steps in the loop extrusion process by individual yeast condensins. The measured median step sizes range between 20-40 nm at forces of 1.0-0.2 pN, respectively, comparable with the holocomplex size. These large steps show that, strikingly, condensin typically reels in DNA in very sizeable amounts with ~200 bp on average per single extrusion step at low force, and occasionally even much larger, exceeding 500 bp per step. Using Molecular Dynamics simulations, we demonstrate that this is due to the structural flexibility of the DNA polymer at these low forces. Using ATP-binding-impaired and ATP-hydrolysis-deficient mutants, we find that ATP binding is the primary step-generating stage underlying DNA loop extrusion. We discuss our findings in terms of a scrunching model where a stepwise DNA loop extrusion is generated by an ATP-binding-induced engagement of the hinge and the globular domain of the SMC complex.

## INTRODUCTION

Structural maintenance of chromosome (SMC) protein complexes, such as condensin, cohesin, and SMC5/6, are vital in many genetic processes, including mitotic chromosome organization and segregation, regulation of sister chromatid pairing, DNA damage repair and replication, and regulation of gene expression (1–6). Emerging evidence points to SMC complexes as universal DNA-extrusion enzymes that play a key role in the macroscale chromosome organization. Chromosome-conformation-capture studies (Hi-C and derivatives) (7, 8) and 3D polymer simulations showed that loop extrusion by condensin can efficiently generate compaction of chromatin fibers (9, 10). Recent *in vitro* single-molecule fluorescence imaging (11–15) demonstrated that SMC complexes extrude DNA into large loops by an ATP-dependent mechanochemical motor activity. These studies showed that the condensin complex constitutes a motor that reels in and extrudes DNA at a very high speed (~600 bp/s) while consuming only very low amounts of ATP (~2 ATP/s) (11), implying that extrusion proceeds in large steps of the order of the ~50 nm condensin size. However, the underlying mechanism of DNA extrusion that leads to the generation of large DNA loops in a step-wise fashion remains to be elucidated.

Structural studies of condensin have suggested various sizeable conformational changes of the protein complex that are potentially associated with the loop-extrusion stepping (16–19). Condensin consists of a ring formed by two 50 nm long antiparallel coiled-coil SMC arms that are mutually engaged by a hinge domain, and, at the opposite site of the ring, a globular domain consisting of two ATPase heads of the SMCs, the Brn1 kleisin, and two HEAT-repeat domains, Ycs4 and Ycg1 (Supplementary Figure 1). A recent AFM study (18) showed large reversible hinge-to-globular domain motions of 22 ± 13 nm. Regarding the mechanochemical ATP hydrolysis cycle that drives the DNA extrusion, this AFM study of yeast condensin as well as a cryo-EM study of cohesin demonstrated that ATP binding induces large conformational changes of the hinge domain from an extended state to a collapsed B shape (18, 20, 21). In attempts to measure condensin-mediated DNA extrusion step sizes, previous single-molecule magnetic tweezers (MT) studies showed significantly varying values, ranging from ~80 nm for *Xenopus* condensin I (22) to ~200 nm for yeast condensin (23) – significantly larger than the 50 nm condensin complex. The large reported step sizes are potentially affected by the intrinsically large thermal motion of the DNA tethers, which renders it very challenging to perform high-resolution step-size measurements at the very low DNA stretching forces where condensin is active (< 1 pN) (11, 23). To elucidate the origin of the large discrepancies in previously reported step sizes, which bears direct relevance for the modelling of the loop-extrusion process, an accurate step size measurement of condensin-induced DNA loop extrusion is much needed.

In this study, we aimed to accurately measure single DNA-loop-extrusion step sizes as well as identify the step-generating process in the ATP mechanochemical cycle using high-throughput single-molecule magnetic tweezers (MT). We optimized the MT using a relatively short 1.5 kbp DNA construct to rigorously resolve small steps with a resolution down to ~10 nm even at the sub-picoNewton forces where condensin is active. We found that individual condensins do extrude loops of DNA in a step-wise fashion, with a force-dependent step size ranging from 17 ± 8 nm to 40 ± 23 nm (median ± median absolute deviation (MAD)) at 1.0 - 0.2 pN DNA stretching forces, respectively. The role of the polymeric nature of the flexible DNA in the loop extrusion process was further probed by Molecular Dynamic simulations of a condensin complex to reel in DNA under low tension. Both the experimental and simulation data revealed that condensin can extrude large amounts of DNA (median ~200 bp), and occasionally even very large amounts (>500 bp), per single extrusion step at low DNA stretching forces. Importantly, the use of an ATP-binding-deficient Q-loop mutant and especially an ATP-hydrolysis-deficient EQ mutant unequivocally demonstrated that ATP binding is the primary step-generating process during DNA extrusion (24, 25), while ATP hydrolysis enables the motor to perform subsequent DNA extrusion step cycles in a consecutive fashion.

## MATERIAL AND METHODS

### Preparation of protein and DNA

*S. cerevisiae* wild type condensin, as well as the EQ (Smc2_E1113Q_-Smc4_E1352Q_) and Q-loop (Smc2_Q147L_-Smc4_Q302L_) mutants, were expressed and purified as previously described (11). Singly biotinylated, linear dsDNA constructs with a length of 1.5, 3.4, and 10 kbp were synthesized *via* PCR and enzymatically ligated to digoxigenin-enriched DNA handles. Primers (Supplementary Table 1) were obtained from Ella Biotech GmbH, Germany or Integrated DNA Technologies, Europe. Biotinylated DNA fragments of different length were produced by using biotin-labeled forward primers (1.5 kb: JT-273; 3.4 kb: TL-34; 10 kb: TL-101) and reverse primers (1.5 kb: JT-275; 3.4 kb: TL-35; 10 kb: TL-102) that contain a *Bsa*I restriction site (Supplementary Table 1) on pBluescript II SK+. To create the digoxigenin (DIG)-enriched handles, a 485 bp fragment from pBluescript II SK+ (Stratagene, Agilent Technologies Inc., USA) was amplified by PCR in the presence of 1:5 Digoxigenin-11-dUTP:dTTP (Jena Bioscience, Germany) using primers CD21 and CD26 (Supplementary Table 1). Prior to ligations of the DNA fragments and handles, all amplicons were digested with the non-palindromic restriction enzyme *Bsa*I-HFv2 (New England Biolabs, UK). The ligation of the DIG-handles and biotinylated DNA fragments was carried out overnight using T4 DNA ligase (New England Biolabs, UK).

### ATPase assay

The ATP hydrolysis rate of WT and EQ mutant condensin complexes was measured using a colorimetric phosphate detection assay (Innova Biosciences) in presence of ATP or ATPγS (Supplementary Figure 7). 50 nM of WT condensin or EQ mutant was incubated with 50 ng/μL λ-DNA (Promega) for 15 min in 40 mM TRIS-HCl pH 7.5, 50 mM NaCl, 2.5 mM MgCl_2_, 2 mM DTT, 5 mM ATP(γS). Afterwards, the concentration of phosphate ions was measured using the accompanied standard protocol of the colorimetric phosphate detection assay of Innova Biosciences.

### Magnetic tweezers

The magnetic tweezers setup used in this study was as described previously (26). Briefly, a pair of vertically aligned permanent neodymium-iron-boron magnets (Webcraft GmbH, Germany) 1 mm apart was used to generate the magnetic field. They were placed on a motorized stage (#M-126.PD2, Physik Instrumente) above the flow cell and allowed red LED light to pass through and illuminate the sample. The transmitted light was collected with a 50x oil-immersion objective (CFI Plan 50XH, Achromat; 50x; NA = 0.9, Nikon), and the bead diffraction patterns were recorded with a 4-megapixel CMOS camera (#Falcon 4M60; Teledyne Dalsa) at 50 Hz. If bead rotation was needed, the magnet pair was rotated with a DC servo step motor (C-150.PD; Physik Instrumente) around the illumination axis. Image processing of the bead diffraction patterns allowed us to track the real-time positions of DNA-bound magnetic beads and surface-bound polystyrene reference beads. The bead *x, y, z* position tracking was achieved using a cross-correlation algorithm in a custom LabView software (2011, National Instruments Corporation) (27, 28). The software also applied spectral corrections to correct for camera blur and aliasing. Up to 300 beads were tracked simultaneously with an approximate Z-tracking resolution of ~2 nm.

### Single-molecule condensin-driven loop-extrusion MT assay

Liquid flow cell preparation and DNA tethering has been described in detail elsewhere (Janissen et al., 2018). Briefly, streptavidin-coated superparamagnetic beads (MyOne DynaBeads, LifeTechnologies, USA) with a diameter of 1 μm were used within this study. Commercially available polystyrene beads (Polysciences GmbH, Germany) with a diameter of 1.5 μm were used as reference beads fixed to the glass surfaces. The polystyrene reference beads were diluted 1:1500 in PBS buffer (pH 7.4; Sigma Aldrich) and then adhered to the KOH-treated (Invitrogen) flow cell glass surface. Afterwards, digoxigenin antibodies (Roche Diagnostics) at a concentration of 0.1 mg/ml in PBS buffer were incubated for ~1 hour within the flow cell, following passivation for ~2 hours of 10 mg/ml BSA (New England Biolabs). After removing non-adhered BSA by washing the flow cell with PBS, 1 pM DNA in PBS was incubated for 30 minutes at room temperature and washed out with PBS. The subsequent addition of 100 μL streptavidin-coated superparamagnetic beads (diluted 1:100 in PBS buffer; MyOne #65601 Dynabeads, Invitrogen/Life Technologies) with a diameter of 1 μm resulted in the attachment of the beads to the biotinylated end of the DNA. Prior to conducting the condensin-mediated loop extrusion experiments, the DNA tethers were assessed by applying a high force (8 pN) and 30 rotations to each direction. Only DNA tethers with singly bound DNA and correct DNA end-to-end lengths were used for the subsequent single-molecule experiments.

To measure the ATP-dependent DNA loop extrusion processes (Figures 1 and 3), we tethered torsionally unconstrained, linear dsDNA of a certain length (1.5, 3.4, and 10 kbp) between magnetic beads and a glass surface (Figure 1A). At a constant applied force of 8 pN, a very low concentration (1 nM) of condensin was injected together with 1 mM ATP into the flow cell, using the same buffer conditions that were previously used to study single condensin-mediated DNA loop extrusion (11). After incubation for 8 minutes, the applied force was rapidly lowered (within 0.8 s) to a force below 1 pN (examples shown for 0.2 pN in Figure 1B and 0.4 pN in Figure 3A and Supplementary Figure 4A) to monitor the stepwise decrease in the DNA end-to-end length.

**Figure 1.**
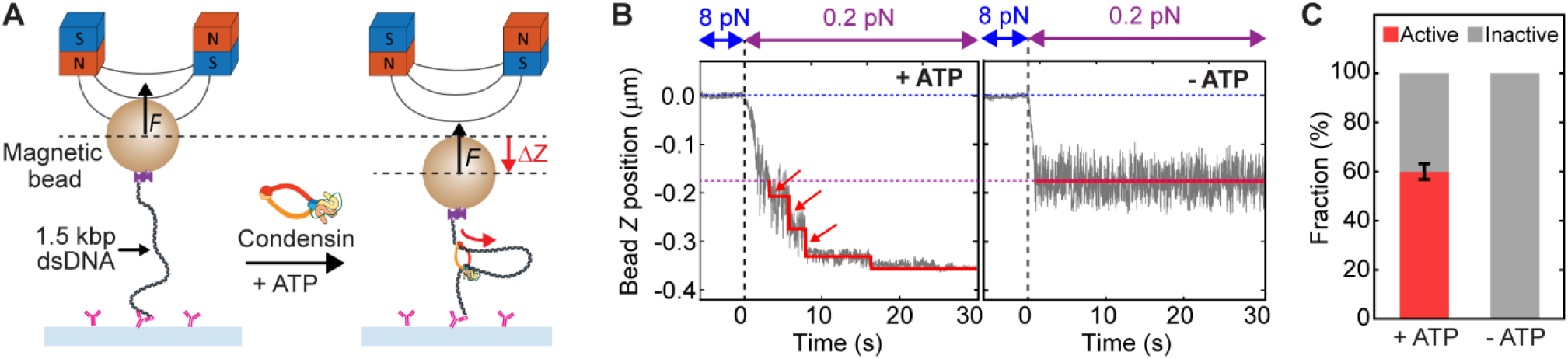
Step-wise DNA loop extrusion by a single condensin holocomplex. (**A**) Schematic of the experimental magnetic tweezers assay to monitor DNA loop extrusion by single condensins. The dotted lines showed the positions of before and upon DNA loop extrusion. (**B**) Representative trajectories of active condensin showing step-wise DNA loop extrusion in the presence of ATP (left), and inactive condensin in absence of ATP (right), at 0.2 pN DNA stretching force. Blue dotted line depicts the DNA tether length at 8 pN, and the magenta dotted line the length of bare DNA at 0.2 pN. Arrows indicate steps in the trace. Red lines are fitted steps from the step-finding algorithm. (**C**) Relative occurrence of active and inactive traces in the presence and absence of ATP for 1 nM condensin (*N* = 207 and 91, respectively). Inactive trajectories represent either bare DNA where no condensin was bound (due to the low concentrations of condensin employed), or DNA where a condensin complex bound but could not perform loop extrusion, e.g., due to the absence of ATP.

For force-titration experiments, dsDNA tethers were re-used for multiple flush-ins of fresh constituents. Previous single-molecule studies did show that DNA-bound condensin after ATP hydrolysis remains bound to the DNA and is still active after high-salt wash for up to 1 hour (23). Condensin was added to the flow cell in reaction buffer (20 mM Tris pH 7.5, 50 mM NaCl, 1 mM ATP, 2.5 mM MgCl_2_, 1 mM DTT, 40 μg/μL BSA) at a concentration of 1 nM while applying 8 pN to the DNA tethers. Afterwards, the force was lowered to 0.4 pN to verify condensin binding and DNA loop-extrusion activity for 90 seconds before the attached bead was able to reach the surface, and the again to 8 pN. To wash unbound condensin out, the flow cells were washed with 500 μl reaction buffer containing 500 mM NaCl and incubated for 5 minutes. After additional washing with 300 μl of reaction buffer, ATP was injected to re-initiate DNA loop extrusion and the force was instantly (within ~1 s) adjusted to the force of interest (0.2, 0.3, 0.4, 0.5, 0.7, 1.0, 2.0, 3.0, 4.0, and 5.0 pN).

After a 5-minute observation time, the force was instantly brought back to 8 pN, accompanied by magnetic bead rotation of 20 times in each direction, to induce full DNA length recovery and rendering the DNA tethers ready for another experiment round. The same process was repeated for various forces, and care was taken to keep the total measuring time within the time condensin stayed active (typically, <40 min).

### Data analysis

The single-molecule data was processed with IGOR Pro, as previously described (Janissen et al., 2018), and further analyzed by custom-written MATLAB scripts. From our raw data, we first removed traces showing surface-adhered magnetic beads and apparent short DNA tethers where the DNA-bead attachment points were far from the magnetic equator of the beads. To do this, the method described in the previous paper (29) was used. Tethers that detached from the surface during the measurement were also rejected from further analysis.

All traces resulting from experiments conducted at identical conditions were pooled by concatenating the traces together into a single time-dependent series. Prior to the step-detection analysis, all traces were filtered using a sliding median average filter over 10 data points to reduce the effect of Brownian noise. An automated step detection algorithm, described in (30), was then applied to the pooled traces for non-biased step detection. Trajectories of condensin-mediated activities were classified according to their behavior, such as loop extrusion steps consisting of consecutive forward steps, single forward steps, and single reverse steps (see examples shown in Figure 4 and Supplementary Figure 4A). The changes in measured bead z-positions for all trajectories were converted to base pairs using base pair length values resulting from prior measured relation between the DNA end-to-end length and the applied force (Supplementary Table 2). Forward step dwell times were measured as the time between two consecutive forward steps, and the reverse step dwell times as the time between a reverse step and the preceding forward step (Supplementary Figure 4B). In the analysis of dwell times, the dwell times of the first step were not taken into account to avoid the inclusion of spurious steps sizes that occurred while still lowering the applied force. In addition, to avoid biasing the results due to the interaction between surface and bead, forward and reverse steps that occurred close to the surface were also not included in our analysis. Summing up, we made our estimate of the step size as conservatively as possible to avoid erroneous results.

### Step validation and detection limit experiments

All step validation experiments were performed with PBS buffer, supplemented with 40 μg/mL BSA. Otherwise, tethered beads were prepared in identical fashion as the experiments involving condensin. Prior to the experiments, a tether test was performed to evaluate the quality of DNA tethers by applying a high force and rotations. Only tethers with singly bound DNA, correct end-to-end lengths, and non-surface adhered magnetic beads were used for the step detection experiments.

In the step validation experiments, the magnetic force was set to a fixed low value (ranging from 0.2 pN to 5 pN). Then, the piezo holding the objective was set to step up or down every 10 seconds, in multiples of 10 nm. Each sequence contained steps from 50 nm down to 10 nm, in multiples of 10 nm. Because the piezo stepping changes the position of the focal plane relative to the bead, each bead exhibited an apparent motion in multiples of 8.4 nm; a difference that stems from the change in refractive index from immersion oil/glass to water. Depending on the pulling force, these steps were more or less submerged within the Brownian motion of the beads. After tracking the thermal fluctuations of the beads, a drift correction was applied by using the average of traces of the reference beads subtracted with the known piezo steps. Next, traces were filtered to 2 Hz with a moving median filter and subjected to step analysis (30). The parameters of the step finding algorithm were kept constant for all stepping experiments. From the step analyses, we obtained a collection of detected steps, each determined by a time of occurrence and a detected step size. This dataset was then compared with the known steps from the piezo motion. A step detection was judged correct if there was a piezo step nearby within 10% (~1s) of the expected dwell time and 30% of the expected step size. Per step size, an average success-of-detection percentage was determined over all traces. Analogously, the reverse was checked as well: a piezo step was deemed ‘found’ if it was seen back in the detected data using the same margins. We found that both types of detection evaluation yielded equal success percentages, as it should be expected for well-tuned step detection. We used the average of these two to obtain for each force a success percentage as a function of step size. The step size where this success percentage crossed 50% was then taken as the detection limit for each force.

To estimate the step detection limit of loop-extrusion steps with different dwell times (between 1 s and 8 s), we performed a series of dwell-time shortening on the 1.5 kbp DNA data presented in Figure 2 by arbitrarily cutting out the plateaus between each piezo step, and concatenated the truncated traces (Supplementary Figure 3A). Using this approach, we effectively shortened expected dwell times in a series to shorten the plateaus from 10 s to 1s, and we determined the step detection limit by the above-described analysis (Supplementary Figure 3B). We observed no significant difference in the step detection performance (Supplementary Figure 3C) for this range of dwell times. We understand this from the design of the used step-finder algorithm, which conservatively limits detection to steps to a step size of order unit noise (31).

**Figure 2.**
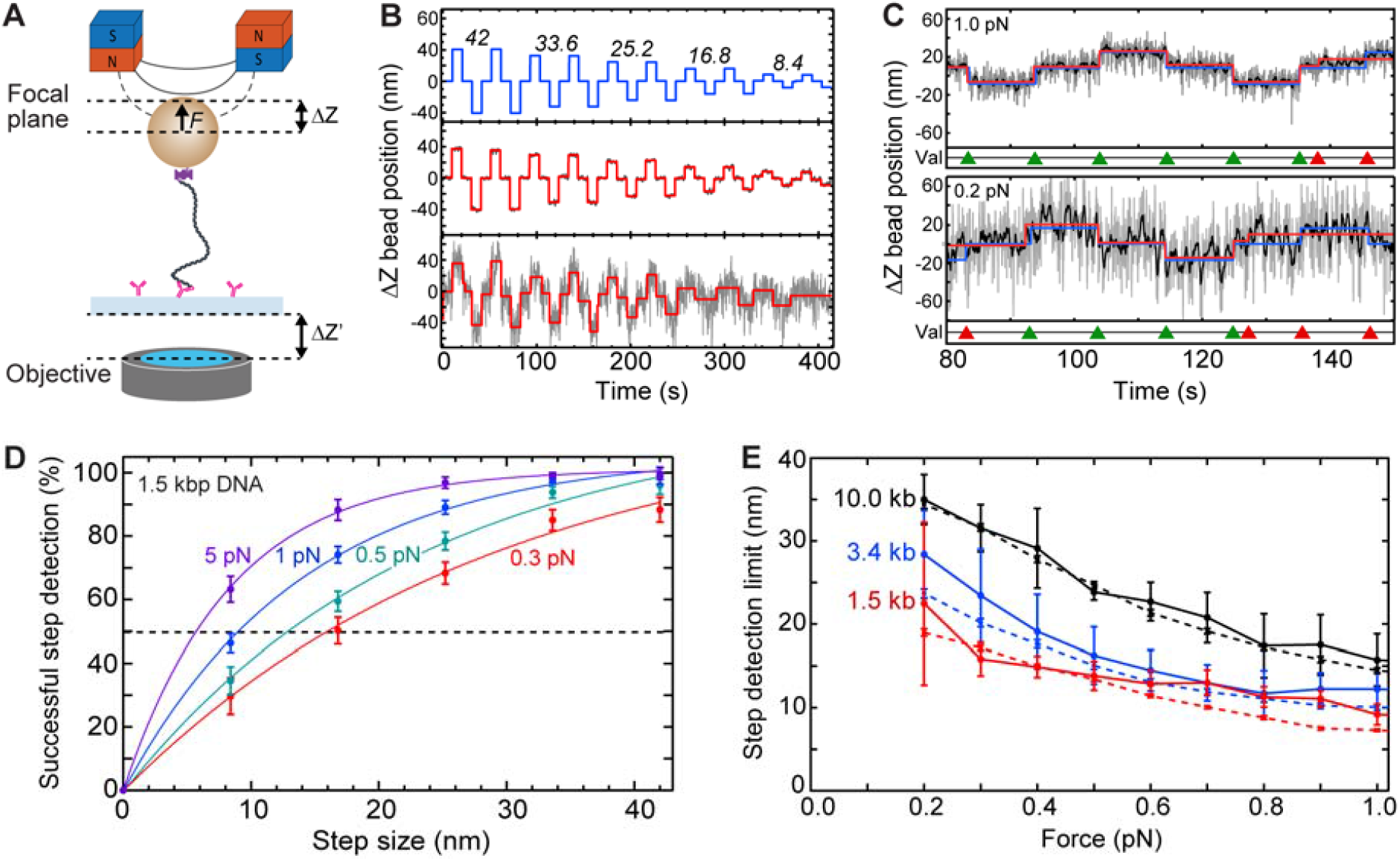
Step validation and step-detection limits for different DNA lengths and stretching forces. (**A**) Schematic for inducing changes in apparent bead Z position by changing the focal plane distance (Δ*Z*) relative to the bead. To do so, the distance between objective and sample surface (Δ*Z*’ = 10, 20, 30, 40, 50 nm) was changed, which in turn changed the focal-plane distance relative to the tethered bead (Δ*Z* = 8.4, 16.8, 25.2, 33.6, 42.0 nm, respectively, accounting for the refractive index mismatch). (**B**) Example trajectories of induced Δ*Z* steps (top; Δ*Z* values in nm depicted above the corresponding induced steps), and corresponding measured bead Δ*Z* positions of a 1.5 kbp DNA at a high (5 pN; center) and low (0.2 pN; bottom) force. Δ*Z* bead position trajectories (grey) were fitted (red) with a step-finding algorithm. (**C**) Zooms of trajectories described in (B) at 1 pN (top) and 0.2 pN (bottom) DNA stretching forces. Δ*Z* bead position trajectories (grey) were filtered (black) to 2 Hz using a moving median filter, prior to applying the step-finding algorithm (red). The induced changes in Δ*Z* position (Δ*Z* = 16.8 nm in the shown example) are superimposed (blue). The step-finding algorithm resulted in successful (green triangles) or unsuccessful (red triangles) detection of induced steps. (**D**) Probabilities for successful step detection (mean ± SD) versus step size for 1.5 kbp DNA at different applied DNA stretching forces. The step-detection resolution is defined as the step size where steps are successfully detected with a 50% probability (dashed line). (**E**) Step detection resolution limit versus DNA stretching force for different DNA lengths (1.5, 3.4, 10.0 kbp DNA; *N* > 80 molecules for each data point). The transverse Δ*Z* bead fluctuations <σ_Z_> (dashed lines) resulting from bead Brownian motion are plotted as well. See also Supplementary Figure 3.

### Molecular dynamic simulation of DNA loop extrusion

We performed Molecular Dynamics simulations of linear 1.5 kbp DNA, modelled as a semi-flexible torsionally unconstrained (as in the experiments) polymer made up of beads of size σ = 10 nm in implicit solvent. DNA beads were held together by finite-extension nonlinear elastic (FENE) bonds, introduced in the equations of motion with the potential

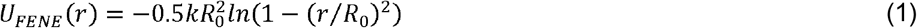

For *r* ≤ *R*_0_ and ∞ otherwise. Here, *r* is the distance between the centers of the bonded beads, *R*_0_ = 1.5 σ is the maximum extension of the bond, and *k* = 30 *ϵ*/*σ*^*2*^ (*ϵ* = *k*_*B*_*T* is the energy scale). The excluded volume interactions between DNA beads were governed by the Weeks-Chandler-Anderson (WCA) potential

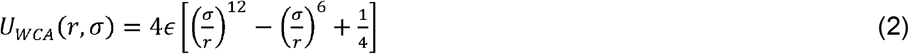

for *r* ≤ *r*_*C*_ and 0 otherwise. This represents a truncated and shifted Lennard-Jones potential with minimum at 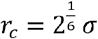, and models purely repulsive interactions. DNA stiffness was accounted for by imposing a bending energy penalty on triplets of neighboring beads, described by the Kratky-Porod potential

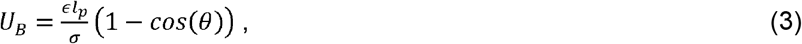

where θ is the angle between consecutive bonds (i.e. segments connecting the centers of two bonded beads) and *l*_*p*_ is the persistence length of the polymer. We set 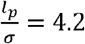, according to the measured *l*_*p*_ = 42 nm (Supplementary Table 2). In the simulated model, both ends of the DNA polymer were tethered, one end to an impenetrable wall, the other end to the surface of a micron bead (see Figure 3I). The micron bead was modelled as a bead of the size of 100 *σ* (corresponding to 1 μm in diameter) and it could only translate vertically along *z*, the direction perpendicular to the horizontal wall. To mimic the experimental assay, we applied an external constant force to the micron bead center, pulling it up along *z*, i.e., away from the wall.

**Figure 3.**
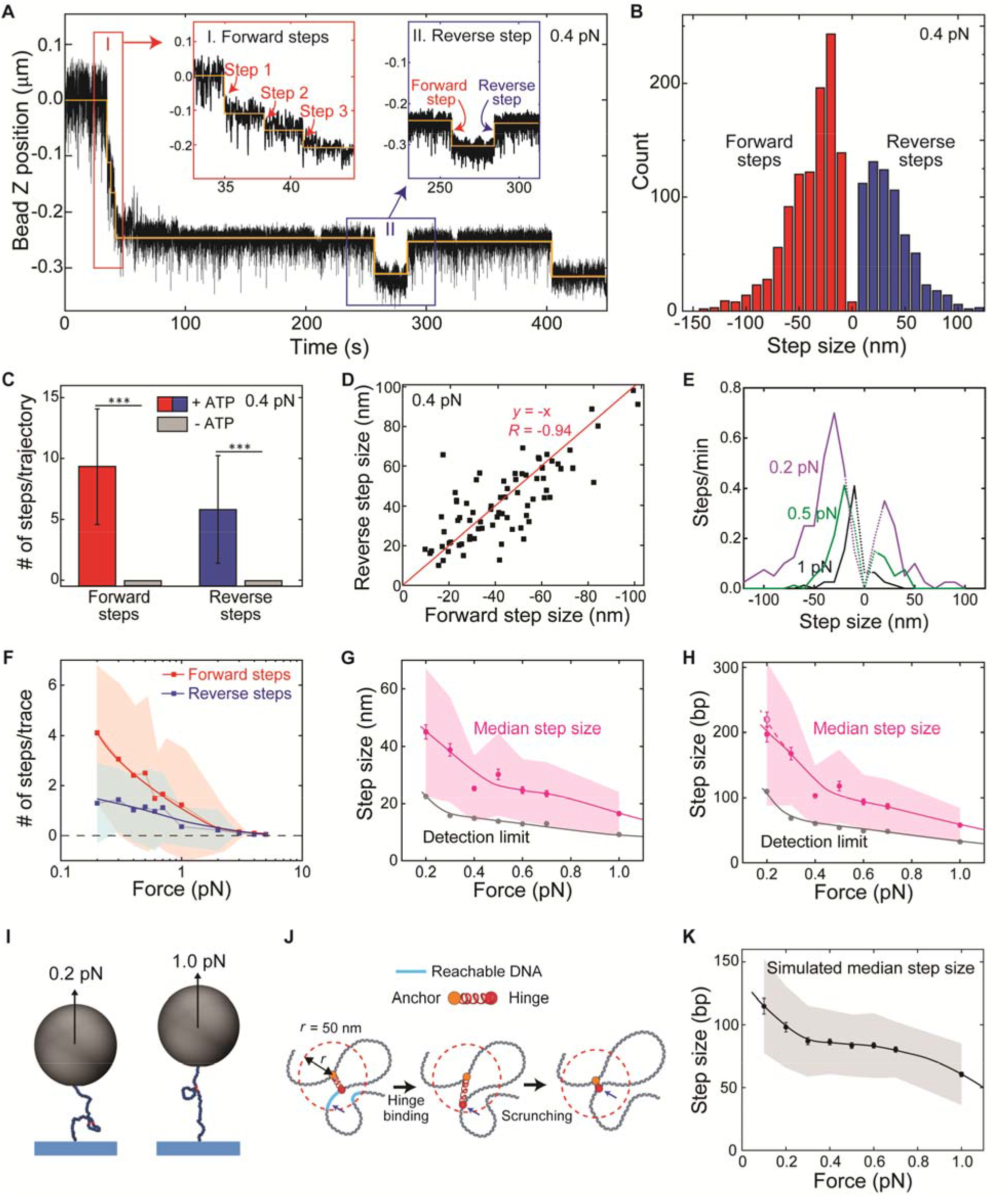
Broad range of force-dependent step sizes of condensin-mediated DNA loop extrusion. (**A**) Representative DNA loop-extrusion trajectory measured at 0.4 pN. Orange lines are fits of the step-finding algorithm. Insets show zooms with consecutive forward steps (left inset) and single forward and reverse steps (right). (**B**) Step size distribution for the 0.4 pN data (*N* = 1,727). Negative step values denote forward steps while positive values are reverse steps. (**C**) Observed number of forward and reverse steps per trajectory of 15 min in the presence and absence of ATP at 0.4 pN (mean ± SD). Statistical analysis consisted of an unpaired two-tailed t-test (*** indicates *p* < 0.001). (**D**) Scatterplot of reverse step sizes versus the corresponding preceding forward step sizes (*N* = 80). Red line depicts a linear fit,y = −xPearson correlation coefficient *R* = −0.94. (**E**) Step size distributions for different DNA stretching forces. Dotted lines denote the range below the step detection limit. Data for other forces are provided in Supplementary Figure 5D. (**F**) Average number of forward and reverse steps (mean ± SD) per 2-minute trace (*N* = 108, 90, 67, 73, 48, 57, 31, 23, 11, 6, 4 from 0.2 to 5.0 pN). (**G**) Combined forward and reverse step sizes (*N* >500 for each force) in nm (median ± SEM; magenta-shaded area corresponds to MAD) verse force for 1.5 kbp DNA constructs. (**H**) Same as G, but step size now given in base pairs. Step size in bp was calculated using the measured relation between the DNA end-to-end length and the applied force (Supplementary Table 2). The detection limit was obtained from Figure 2E and Supplementary Figure 3. Solid magenta and grey lines in (G) and (H) represent guides to the eye. (**I**) Simulated DNA topology at 0.2 pN and 1.0 pN with a loop extruded by condensin. **(J)** Schematic of the methodology used in MD simulations. Condensin is modelled as a spring bringing together two DNA segments. The position along DNA of one of its ends (hinge, red) is periodically updated: reachable DNA (cyan) lies within 50nm from the other, fixed, end (anchor, orange). Step lengths were computed from the average height of the simulated micron bead in between consecutive steps. **(K)** Simulated median step sizes in bp versus force for 1.5 kbp DNA constructs (*N* = 213, 291, 305, 329, 328, 355, 353, 409 from 0.1 to 1.0 pN).

Each simulated DNA polymer was loaded with a condensin, which was modelled as a spring bringing together two DNA beads. The spring was described by a harmonic potential

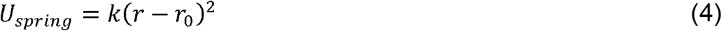

Where *k* = 4*k*_*B*_ *T*/*σ*^2^ and *r*_0_ = 1.6 *σ* = 16 nm are, respectively, the spring stiffness and resting distance (we chose the resting distance comparable with the minimum size of condensin globular domain). To model loop extrusion, we adopted a simpler version of the framework proposed by Bonato *et al*., 2021 (32). Condensin is initially loaded at a random position along the polymer and forms a short polymer loop by connecting two beads (*i* and *i*+2). At the time of the loading, one of the ends of the spring was labelled as anchor, the other as hinge, as that is presumably the second binding site in the motor that derives the DNA loop extrusion. As the simulation progressed, condensin extruded progressively larger loops as follows (see Figure 3J). While the bead position of the anchor was kept strictly fixed (similar to the anchoring Ycg1 protein in the condensin holocomplex), the position of the hinge was periodically updated. Every time the position changed, a new position was selected by randomly picking one polymer bead Within 5 *σ* = 5.0 nm (Euclidean distance) from the head at the time of the update (this value was chosen as compatible with the maximum size of the complex). To avoid the condensin taking steps back or undoing the extruded loop, we limited the selection of the new position to the beads that were (i) on the side of the hinge and (ii) outside the extruded loop (see Figure 3J). Note that in this way the extrusion direction was fixed. The extrusion proceeded until the hinge met either the surface wall or the micron bead.

For each different pulling force, we simulated 40 polymer chains. We started by equilibrating the system for a sufficiently long time (3×10^7^ integration time steps) before we loaded the condensin and subsequently ran the simulation for 2×10^9^ time steps. The integration time step was set to 0.01τ_B_, where τ_B_ was the time it takes a particle of the size σ to diffuse by its size; τ_B_ = 2.2×10^−6^ s for a 10 nm particle in water. The position of the hinge of a condensin was updated every 10^8^ time steps, i.e. every 2.2 s. The equations of motion of the beads, accounting for the implicit solvent (Langevin heat bath) and the potentials described above, were integrated using the LAMMPS package (https://lammps.sandia.gov/). In practice, the update of the condensin to simulate extrusion was done by calling LAMMPS as a library from a C++ program.

## RESULTS

### Magnetic tweezers resolve single DNA extrusion steps of individual condensins

In order to resolve single DNA-loop-extrusion steps that are induced by an individual *S. cerevisiae* condensin holocomplex in real-time, we employed a single-molecule assay based on MT (33) which is a particular advantageous technique because it allows to observe real-time changes in the end-to-end length of a single DNA molecule at high spatiotemporal resolution (34). Here, we optimized the MT technique by using a short DNA construct (1.5 kbp) to minimize the Brownian motion to its limits such that we could detect very small steps down to a resolution of ~10 nm even at sub-pN forces. Using this technique, we were able to measure up to 300 DNA tethers simultaneously at a frame rate of 50 Hz, which provides large data sets suited for statistically robust analysis (23, 26, 35).

We monitored the asymmetric condensin-mediated loop-extrusion process on torsionally unconstrained 1.5 kbp linear dsDNA (Figure 1A) in the presence of 1 nM condensin and 1 mM ATP, i.e. the same conditions previously used to study single condensin-mediated DNA loop extrusion (11). After a sudden force change from 8 pN to a low force (i.e. 0.2 pN in Fig.2A), we observed a stepwise decrease in the DNA end-to-end length, which can be attributed to condensin-driven DNA loop extrusion activity (Figure 1B; left). Due to the arbitrary binding position of the condensin onto the extended DNA molecule (11), we observed different final degrees of DNA compaction, with ~50% on average (Supplementary Figure 2). Importantly, in the absence of ATP, changes in DNA lengths were never observed (Figure 1B; right), confirming that the reduction in DNA lengths in experiments with ATP resulted from an ATP-dependent DNA-loop-extrusion processes (Figure 1B; left), in agreement with previous studies (22, 23). Counting the fraction of DNA tethers that exhibited a stepwise reduction in apparent DNA length (Figure 1C), we found that ~60% of the tethers showed such loop extrusion activity at this low 1 nM concentration. This activity can be largely attributed to a single condensin, since the fraction of multiple condensin acting on a single DNA strand is very low, in agreement with previous MT studies that showed non-cooperative condensin-driven DNA compaction for condensin concentrations between 1 and 10 nM (23).

### Magnetic tweezers can be optimized to resolve 10-20 nm step sizes at sub-pN forces

We determined the minimally resolvable step size for different DNA tether lengths at forces ranging from 0.2-5 pN. The resolvable step size is largely limited by the intrinsic noise in MT measurements due to force-dependent bead fluctuations resulting from Brownian motion (36). To quantitatively assess the resolution of our assay, we induced artificial steps of different sizes (Δ*Z* = 8.4, 16.8, 25.2, 33.6, 42.0 nm) by changing the focal plane relative to the DNA-tethered magnetic bead (Figure 2A), and subsequently determining to what extent we could resolve these user-induced steps. The example traces shown in Figure 2B show that, as expected, the observable Brownian noise reduced at higher forces. To assess to what degree we can successfully re-detect the induced steps (Figure 2B: top), we fitted a step-finding algorithm (30) to our data using Chi-squared minimization without any fit parameter boundaries with respect to step sizes or locations. The example traces in Figure 2B (middle) show that at high force (5 pN) the smallest induced step size of Δ*Z* = 8.4 nm could readily be detected, while at the lowest applied force (0.2 pN; Figure 2B: bottom), the smallest steps were outweighted by the noise.

In order to determine the minimal detectable step size at different forces, we applied a validation algorithm that compared the detected steps by the step-finding algorithm with the artificially induced steps (Figure 2C; red lines and blue lines, respectively). We defined a step relocation as ‘successful’, if both the step size and the dwell time were close to the user-induced values. The example traces shown in Figure 2C for 1.0 (top) and 0.2 pN (bottom) show the comparison for induced steps of Δ*Z* = 16.8 nm. Successful relocations are indicated with green triangles, whereas red triangles denote false positives and false negatives, i.e., the allocation of steps when no steps were induced, or induced steps that remained undetected, respectively. Over a range of step sizes and forces, these validations were repeated for ~100 traces with 40 induced steps at each condition. From the results, shown in Figure 2D, we observed that lower forces led to a lower step re-detection success, as expected. Since proper step-detection should avoid both under-fitting and over-fitting, we defined the step-detection limit as the step size where the false positives and false negatives together average to 50% (Figure 2D: dotted line). We found that the step detection limit decreased from 16 to 6 nm for applied forces that increased from 0.3 pN to 5 pN, respectively.

Figure 2E displays the detection limit as a function of force for various DNA lengths. The obtained step detection limits closely followed the trend of the bead’s Brownian noise in the z position over the entire tested force range (Figure 2E and Supplementary Figure 3A: dashed lines). This indicates that Brownian noise is the dominant parameter that determines the step detection limit. The dwell time duration did not change the detection limit (Supplementary Figure 3B). Longer DNA tethers with lengths of 3.4 and 10 kbp exhibited higher step detection limits (3.4 kbp: ~7 to ~30 nm; 10 kbp: ~7 to ~35 nm) than the 1.5 kbp DNA tethers, throughout the entire force range. Importantly, at low forces <1 pN where condensin is active, the low detection limits for the 1.5 kbp DNA construct (~10 to ~20 nm between 1.0 to 0.2 pN, respectively) are well below the ~50 nm size of the condensin holocomplex and thus below the putative step size in loop extrusion. In subsequent DNA-loop-extrusion experiments, we used the 1.5 kbp DNA construct for measuring step sizes.

### Condensin-induced DNA loop extrusion exhibits a broad distribution of step sizes

With the demonstrated ability to resolve small step sizes, we next characterized the steps during DNA loop extrusion induced by condensin under various conditions in more detail. Figure 3A and Supplementary Figure 4A show typical 1.5 kbp DNA-compaction traces, where two distinct signatures are observed: a series of ≥2 consecutive forward steps (Figure 3A; left inset), which was the generic DNA loop-extrusion behavior, as well as occasionally a forward step that was followed by a single reverse step upwards (Figure 3A; right inset). Such reverse steps occurred exclusively after a prior forward step. DNA steps occurred almost exclusively very fast, i.e., within a single 20 ms imaging frame.

Employing our step-detection algorithm to the experimental data, we identified the distribution of both forward and reverse step sizes, as exemplified in Figure 3B for 0.4 pN. Notably, the step size distribution was found to be very broad, covering a wide range of small and large step sizes. In the absence of ATP, no step signatures were observed (Figure 3C). The measured forward and reverse dwell times, defined by the time between either two consecutive forward steps or between the reverse step and the preceding forward step, respectively (Supplementary Figure 4B), both followed a single exponential distribution, suggesting that the steps originated from a first-order rate-limiting process. Interestingly, the median step size was found to be comparable for forward and reverse steps, i.e., 32 ± 21 nm for forward steps compared to the 31 ± 23 nm for reverse steps (Figure 3B). The size of individual forward/reverse step pairs was also found to be highly correlated (Figure 3D), i.e. large reverse steps correlated with large preceding forward steps, and small reverse steps with small preceding forward steps, which indicates that the reverse steps are mechanistically linked with the preceding forward steps.

The key result of Figure 3B is the observed median step size value of ~32 nm for the 1.5 kbp DNA length for 0.4 pN, which is twice higher than the step detection limit of ~15 nm (Figure 2E and Supplementary Figure 4C) at this 0.4 pN force. To further validate our result, we compared the experimental data with the step sizes deduced for the user-induced 33.6 nm steps (Supplementary Figure 4D), which are close in value to the 32 nm median value in the condensin experiments. For this user-set step size, our validation test resulted in only a small spread of ~4 nm (MAD), indicating that tracking and step-finding analysis errors were not predominantly causing the broad distribution observed for the condensin stepping (Supplementary Figure 4E). By contrast, the median step sizes resulting from 3.4 and 10 kbp DNA constructs were found to be substantially larger, particularly at forces ≤ 0.4 pN (Supplementary Figure 5A–C). For these DNA lengths the step detection limits (Figure 3E), especially at 0.2 pN with ~30 nm and ~35 nm for 3.4 and 10 kbp DNA, respectively, became similar to the median step size of ~30 nm measured for 1.5 kbp DNA. Notably, we also observed a significant difference between step sizes for forward and reverse steps for the 3.4 and 10 kbp DNA lengths but not for 1.5 kbp DNA (Supplementary Figure 5A–C and Supplementary Figure 5EF), further indicating that the step detection became unreliable for the longer DNA molecules. For these reasons, we limited the analysis to the 1.5 kbp DNA, for a reliable measure of the DNA loop-extrusion step sizes.

### Condensin extrudes DNA in steps of tens of nm, reeling in hundreds of base pairs per step

We observed a pronounced force dependence of the loop-extrusion stepping behavior which reduced with increasing force (Figure 3E,F). Above 1 pN, we could not observe any significant loop extrusion activity, in agreement with previous observations (11, 23). Figure 3E shows the distribution of observed step size from 0.2 pN to 1 pN, which exhibited a distinctly narrower distribution at higher forces. The ratio between reverse and forward steps was largely maintained at a value of about 0.3 at all tested forces (Figure 3F and Supplementary Figure 4F).

A major result of this work is displayed in Figure 3G, which shows that the step sizes increase with lowering the DNA stretching force. Step size values ranged for the 1.5 kp DNA construct from 17 ± 8 to 40 ± 23 nm (MAD) between 1 and 0.2 pN, respectively. The measured values were at all forces significantly larger than the step detection limits (grey line). The width of the step size distributions decreased from 23 nm (MAD at 0.2 pN) to 8 nm (MAD at 1.0 pN) upon application of higher force (Supplementary Figure 4G).

Upon conversion of the measured step sizes in nm to force-dependent DNA steps measured in bp using a measured relationship between DNA end-to-end length and force (Supplementary Table 2), we found that the steps of extruded length of DNA also exhibited a notable force dependence, yielding median lengths of ~60 bp up to ~220 bp between 1 and 0.2 pN per loop extrusion step, respectively (Figure 3H, dashed line). Intriguingly, the *maximum* step size amounted to ~500 bp on average across the entire tested force range, and reaching ~800 bp at the lowest force of 0.2 pN (Supplementary Figure 4H). The observed force dependence and the large extruded DNA lengths suggest that in each step the loop extrusion proceeds when the condensin holocomplex grabs new DNA from within the random polymer blob that DNA forms at low forces.

To more firmly establish that the semi-flexible polymeric nature of the DNA is a key determinant of the force-dependent DNA step sizes, we performed Molecular Dynamics simulations (MD) based on a generalized version of the well-known loop-extrusion model that can perform extrusion by grabbing segments of DNA that are non-contiguous (32). In the simulated model (see Methods), one end of a 1.5 kbp DNA was tethered to an impenetrable wall mimicking the surface, and the other end to a 1 μm-sized bead that was free to move in the direction perpendicular to the surface. The simulated conformation of the tethered DNA is illustrated in Figure 3I for 0.2 and 1 pN. DNA was loaded with a simplified condensin motor that was modelled as a spring bridging two DNA segments, where the position of one end of the spring (head) was fixed while the position of the other end (hinge) was periodically updated during the progression of the loop-extrusion simulation (Figure 3J). At each DNA loop-extrusion step, a new hinge position along the DNA was randomly picked within a radius of 50 nm (mimicking the SMC coiled coils size) from the head position.

The simulation showed a step-wise shortening of the end-to-end length (Supplementary Figure 6A) until the bead reached either the surface wall or the bead. Upon analyzing the step sizes from these traces, a clear force-dependence of the step sizes was found (Figure 3K, Supplementary Figure 6B), in agreement with our experimental findings (Figure 3A,H). A median step size of ~60 bp to ~110 bp was observed for forces of 1 pN down to 0.2 pN, respectively. Occasionally, even much larger steps were observed, up to ~550 bp can occur at the low forces (Supplementary Figures 6D), which are close to sizes encountered experimentally. The experimental and simulated median step sizes were largely comparable at forces >0.3 pN, while at lower pulling forces, the simulated step sizes were notably smaller than the experimentally measured values (compare Figure 3H and 3K). To assess a potential over-fitting of the experimental data by the step-finder algorithm, the MD simulation provided the practical means to quantify any existing step detection error. By applying the step-finding algorithm to simulated loop-extrusion traces, we found a systematic overestimate of +10% in the determination of step sizes at 0.2 pN, but without deviations at the higher forces (Supplementary Figure 6C). Hence, we corrected the experimental step size value at 0.2 pN (compare solid versus dashed line in Fig.3H). Notably, the simulation value of ~110 bp at 0.2 pN still deviated significantly from the corrected experimental median step size value of ~200 bp. This quantitative difference suggests that future modelling work on SMC loop extrusion needs to account for additional effects such as enthalpic penalty for grabbing and bending short DNA segments compared to longer ones (37) or other potential non-specific condensin-DNA interactions (38).

Overall, the MD simulations confirm that the semi-flexible polymeric nature of the DNA is essential to understand the force-dependent DNA step size in DNA loop extrusion by condensin. Since our model of loop-extrusion is not explicitly dependent on the external pulling force, we conclude that the force-dependence observed in the step sizes derives from the availability of DNA in proximity of the condensin complex. Our combined experimental and simulation data thus support the idea that the mechanism underlying the stepping in DNA loop-extrusion involves a random grabbing of nearby DNA within the spatial reach of the condensin holocomplex.

### ATP binding is the step-generating stage in the ATP hydrolysis cycle that underlies condensin-mediated DNA loop extrusion

While it is established that condensin extrudes DNA loops in an ATP-hydrolysis-dependent manner (11, 23), our MT experiments interestingly allow to discriminate between the mechanochemical functions of ATP binding and ATP hydrolysis through the study of ATP-mutants of condensin. Specifically, we studied the DNA extrusion stepping behavior of the EQ mutant (Smc2_E1113Q_-Smc4_E1352Q_) that allows ATP binding but blocks its hydrolysis by a Walker B mutation in the ATPase domains (Supplementary Figure 6), as well as for the Q-loop mutant (Smc2_Q147L_-Smc4_Q302L_), which blocks ATP binding to the ATPase domain entirely. The relative occurrence of the different step signatures of observed activity (i.e. consecutive forward steps, single forward steps, single forward followed by reverse step pairs, and inactive traces) is shown in Figure 4E. Contrary to the wild-type condensin (Figure 4A,E), the EQ mutant was *not* able to perform consecutive DNA extrusion steps, as >95% of the stepping traces showed only a single step (and the rare multiple step traces may be attributed to the action of multiple condensins). Instead, the EQ mutant was solely capable of performing single forward steps (Figure 4B,E), which was sometimes followed by a reverse step (Figure 4C,E). The Q-loop mutant did not show any step activity at all (Figure 4DE). In the absence of ATP, all condensin variants lacked step activity. The results clearly demonstrate the necessity of ATP hydrolysis to perform consecutive DNA extrusion steps, while ATP binding (without subsequent hydrolysis) only allows to perform a single step.

**Figure 4.**
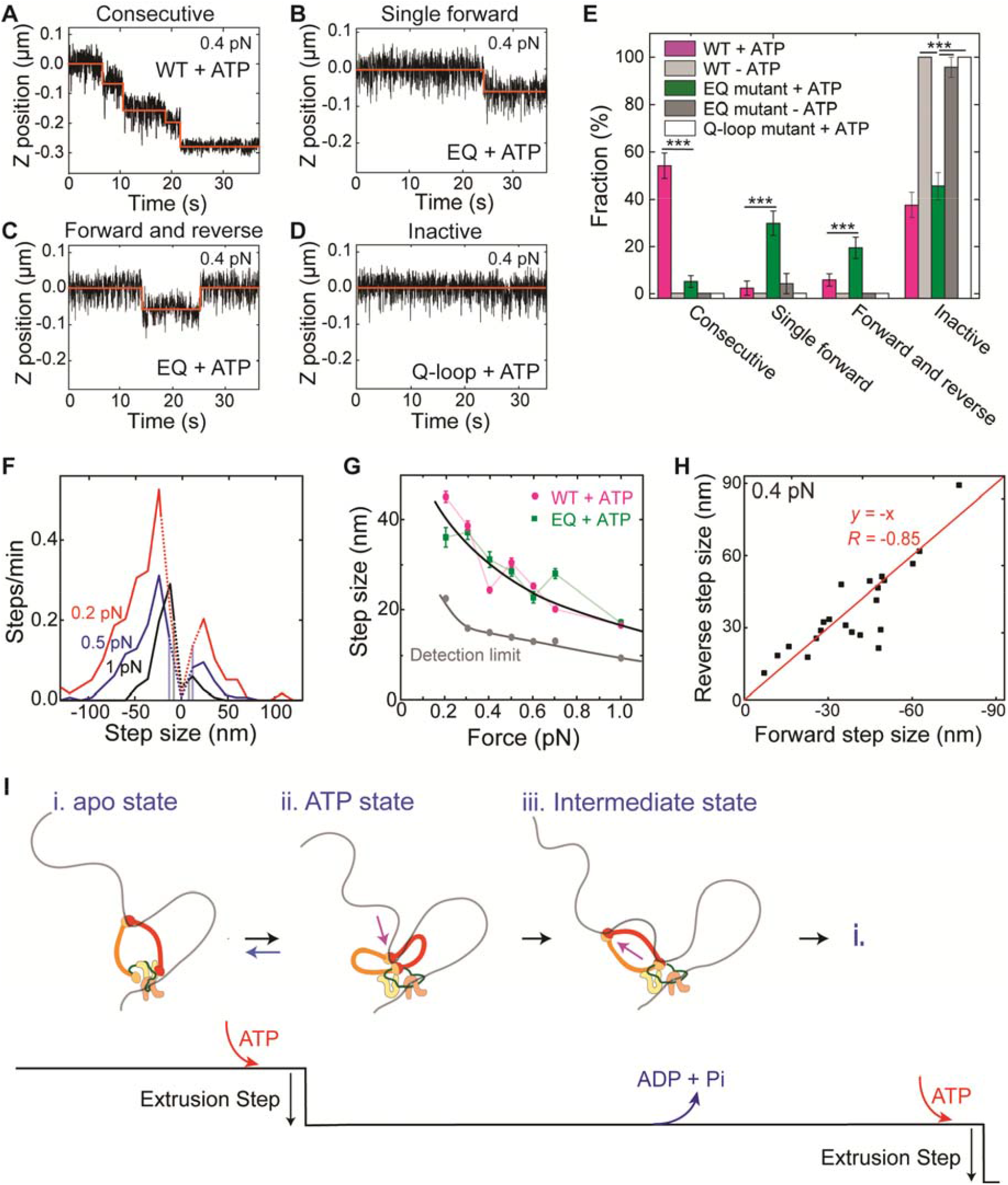
ATP-binding of condensin induces a single step in DNA loop extrusion. (**A–D**) Representative trajectories for (A) consecutive DNA loop extrusion steps, (B) a single forward step, (C) a single forward followed by a reverse step, and (D) an inactive trajectory, all probed in the presence of ATP. (**E**) Observed fractions of different stepping behavior (*N* = 85, 96, 77, and 18 for wild-type with ATP, without ATP, EQ mutant with ATP, and Q-loop mutant with ATP, respectively). Statistical analysis consisted of an unpaired two-tailed t-test (*** indicates *p* < 0.001). (**F**) Step size distributions of the EQ mutant in the presence of ATP at different DNA stretching forces (0.2 pN: N = 154; 0.5 pN: N=186; 1 pN: N=106). Dotted lines denote step sizes below the step resolution limit. Data for other forces are provided in SI. (**G**) Step size in nm (median ± SEM) for WT (magenta) and EQ mutant (green) versus applied force (0.2 pN: *N* = 69; 0.3 pN: *N* = 53; 0.4 pN: *N* = 73; 0.5 pN: *N* = 74; 0.6 pN: *N* = 45; 0.7 pN: *N* = 58; 1 pN: *N* = 76). Forward and reverse steps were pooled together in these data. Solid black and grey lines represent guides to the eye. (**H**) Scatterplot of reverse step sizes versus the corresponding preceding forward step sizes (*N* = 15) for the EQ mutant at 0.4 pN. Red line depicts a linear fit,y = −x Pearson correlation coefficient *R* = −0.85. (*N* = 22) (**I**) Proposed working model for the condensin conformational changes during the ATP hydrolysis cycle. (i) Condensin holocomplex is anchored to DNA by the Ycg1-Brn1 subunit. In the open configuration, the hinge domain binds DNA. Note that the hinge grabs an arbitrary nearby location within the randomly structured DNA polymer coil. (ii) Upon ATP binding to condensin, the SMC ring changes from the open to a collapsed (butterfly) configuration, generating a single DNA loop-extrusion step where DNA is reeled in via the hinge movement. From our data, we conclude that this step is in principle reversible, whereby state *ii* can return to state *i*. (iii) After DNA transfer to the globular domain of condensin, the hinge is released, presumably during ATP hydrolysis, whereupon it is available to bind new DNA for the next step in the cycle. Upon repetition of this cycle, DNA is extruded into an expanding loop in consecutive steps.

The distribution of step sizes as well as the force dependence of the step size for the EQ mutant (Figure 4FG) was very similar to the WT condensin. In addition, we again observed a high correlation between the forward and the reverse step sizes (Figure 4H) for the EQ mutant in presence of ATP, suggesting that the reverse steps may originate from spontaneous release events between the hinge and the HEAT globular domains. The data indicate that the ATP-binding-dependent step for the EQ mutant is identical to the WT step formation, and establish that ATP binding is the primary step-generating process within the ATP hydrolysis cycle.

## DISCUSSION

Using MT with short 1.5 kbp DNA tethers that allow the detection of DNA loop extrusion steps as small as 10–20 nm in the relevant low-force regime, we were able to resolve that condensin extrudes DNA in a stepwise fashion, with a median step size of the order of 20–40 nm (60–200 bp) at DNA stretching forces from 1 to 0.2 pN, respectively. In addition to the observed force dependence of step sizes, we identified that ATP binding is the primary step-generating process during DNA loop extrusion. Below, we discuss several of the most salient findings.

### Step sizes indicate large conformational changes of condensin during loop extrusion

The measured median step sizes ranged between 20 and 40 nm over the relevant range of forces where condensin is active, i.e. only slightly smaller than the longest dimension of the condensin holocomplex with a SMC coiled coils length of about 50 nm. The observed large step sizes define SMCs as an entirely distinct class of DNA-processing motor proteins that are unlike any other DNA-processing enzymes reported before (e.g. helicase, translocases, polymerases), which translocate in single-base pair steps upon each ATP hydrolysis cycle (39–44).

Importantly, our accuracy in measuring DNA loop-extrusion step sizes was significantly better than the step detection limit, which was not quantitatively estimated in previous studies (22, 23). Our measurements of the DNA loop-extrusion step size yielded, for our most reliable data with the 1.5 kbp DNA construct, significantly smaller values than previous estimates of ~80 nm (at 0.4 pN and 2 nM condensin, measured with 4 kbp DNA) and ~200 nm (0.75 pN, 8.6 nM condensin, 20 kbp DNA) (23). As the Brownian noise increases with the DNA tether length (Figure 2E), the small DNA loop extrusion steps may have remained undetected in these studies. Indeed, the larger step sizes and discrepancy between forward and reverse step sizes that we ourselves observed for the 3.4 and 10 kbp DNA lengths (Supplementary Figure 5A–C) strongly support the possibility of bias in observed step sizes when using longer DNA constructs. In addition, higher condensin concentrations can readily increase the propensity of multiple condensins acting on a single DNA tether. Considering that the distribution of single condensin-mediated step sizes is very broad (Figure 3B,E), Brownian noise in the earlier studies may have effectively cut off the smaller step sizes, yielding an average step size that exceeds the intrinsic value. Our observed range of average step sizes (~20–40 nm), is, however, in good agreement with the recently reported hinge-to-globular domain distance of 22 ± 10 nm that is involved in a dynamic toggling between O to B shapes, i.e., from an extended state to a hinge-engaged collapsed state (Supplementary Figure 1) (18). The large DNA step sizes of ~200 bp at low force is similar but somewhat lower than values that can be estimated by combining the loop extrusion speed and ATPase rates in previous *in vitro* studies on condensin and cohesin with ~0.5–1 kbp/s and 2 ATP/s (11, 13, 45), and of SMC-ScpAB from *B. Subtilis* and *C. Crescentus* with 300–800 bp/s and 1 ATP/s (46–48).

### Large DNA extrusion steps are associated with the structural flexibility of DNA at low forces

A broad distribution of condensin-mediated DNA loop extrusion step sizes was observed (Figure 3B,E), with a variation in step sizes that was well beyond that set by the instrumentation, as can be seen from the much narrower variation in the user-induced step sizes in our step-validation tests (Supplementary Figure 3E). One contribution to the variation in measured DNA extrusion step sizes could be a variation in the degree of internal conformational changes of the condensin, similar to a sizeable variation (MAD = 10 nm) in the hinge-to-head motion that was observed previously with AFM (18). Much more importantly, however, is the variation that is induced by the structurally very flexible and dynamic nature of DNA at these very low stretching forces. We suggest that such steps result from the hinge domain grabbing proximate DNA during every DNA loop-extrusion cycle. Indeed, our current as well as recently published model simulations indicate that this scenario can result in large step sizes (37).

Our experimental observations and MD simulations thus support the notion that condensin is able to bind to proximate DNA within the polymer blob of largely unstretched, flexible, and dynamic DNA, to subsequently reel it in (Figure 3H,K and Supplementary Figure 6D). This mechanistic process is entirely different from the characteristics of motor proteins such as myosin and kinesin (49–51), which walk along stiff actin or microtubule protein filaments (52). The very large step size of condensin (20–40nm) is also consistent with the very low stall force of ~0.5 pN of this motor, as this combination may involve ~4 k_B_T of work which can readily be provided by the free energy generated by hydrolysis of 1 ATP per step.

### ATP binding and hydrolysis relate to two distinct mechanistic processes

Our results unequivocally demonstrate that ATP *binding* is the process associated with the DNA-extrusion stepping, since the ATP hydrolysis-deficient EQ mutant (Figure 4E) was able to make single steps, but unable to perform consecutive DNA extrusion steps, while the ATP-binding-deficient Q-loop mutant did not show any DNA extrusion activity. Consequently, and different from most other ATPase motor proteins, ATP hydrolysis occurs downstream in the hydrolysis cycle to enable DNA extrusion steps in a consecutive manner.

Our finding that ATP binding is the step-generation process differs from a previously suggested DNA pumping model (53) that attributed ATP hydrolysis to the generation of a step, associated with the zipping-up of the SMC arms from an ATP-bound O shape into an ATP-unbound I shape that pushes DNA from the hinge to the head domains. Indeed, the I-to-folded state transition was suggested to be triggered by ATP hydrolysis (17, 54). By contrast, our result that ATP binding is the step-generating process is in good agreement with cryo-EM results on cohesin (20, 21, 55, 56) and with AFM data on condensin that showed a transition from an extended state to a hinge-engaged state upon ATP binding (18). In addition, our results are also consistent with recent AFM and single-molecule FRET study on human cohesin that showed that ATP-binding promotes head domain engagement that provides DNA binding sites via hinge-head interaction (57). Importantly, our finding further agrees with results found *in vivo*, which reported that the hinge domain of cohesin engages with the globular domain to form a B shape (58).

Our data are also in good agreement with a DNA scrunching model that we recently hypothesized based on these AFM data (18), where condensin anchors itself to DNA using the safety belt of the Ycg1-Brn1 domains (Figure 4I i) (11, 59), whereupon the hinge domain binds to a proximate region of the DNA, and ATP binding induces a transition from an extended O shape to a collapsed hinge-engaged B shape, thereby pulling the hinge-bound DNA to the globular domain (Figure 4I ii), which establishes a step in the loop extrusion process. In support of this concept, previous studies showed that ATP binding induces the dimerization of the head domains, forming a positively charged cavity that is able to bind DNA (21, 24, 57, 60, 61). The low stalling force that we observed is consistent with a type of a motor mechanism that involves a Brownian ratchet motion of the flexible SMC arms that underlies the hinge to globular domain step (62). In the next stage of the cycle, ATP hydrolysis occurs whereupon the hinge is released and the condensin returns to the O shape where it is available to bind to a new DNA target site for the next step (Figure 4I iii), thus closing the DNA loop extrusion cycle. Additional support for this scenario is provided by a prior study that showed that ATP hydrolysis induces the dissociation of the head dimers, hence disrupting the capability to bind DNA (24). The model indicates that condensin may employ ‘credit-card energetics’ (63) where the energetic costs of compacting the DNA is “spent” by ATP binding prior to the energy “payment” by ATP hydrolysis.

## CONCLUSION

In conclusion, our experimental and MD simulation results demonstrate that the SMC proteins are unique protein complexes that can extrude DNA loops with sizes of hundreds of base pairs in a single step. Our study shows that DNA loop extrusion steps consist of two distinct processes: ATP binding provides the step-generating process where likely DNA bound to the hinge domains is pulled to the globular domains, leading to DNA loop extrusion. In the second subsequent step, ATP hydrolysis presumably enables the hinge domain to target a next DNA site for a subsequent loop extrusion step. The observed strong dependence of step size on the applied DNA stretching force revealed that the flexible nature of the DNA polymer at very low stretching forces facilitates the extrusion of large amounts of DNA in each step in the loop extrusion process, indicating a mechanism that differs from previous motor proteins as well as that of popular models of DNA loop extrusion, as it appears to require to grab non-contiguous DNA segments. Our work reveals unique characteristics of the motor action of condensin which may be conserved among other SMC proteins that exhibit DNA loop extrusion.

## DATA AVAILABLITY

All data that support the findings of this study are available on request from the corresponding author, CD.

## SUPPLEMENTARY DATA

Supplementary Data are available at NAR Online.

## ACKNOWLEDGMENT

We thank J. van der Torre for DNA construct synthesis, Eli van der Sluis for protein purification, and Roman Barth, Allard J. Katan, Eugene Kim, Biswajit Pradhan for discussions. We acknowledge Christian Haering for providing the condensin mutants.

## SUPPLEMENTARY DATA

### SUPPLEMENTARY FIGURES

**Supplementary Figure 1.**
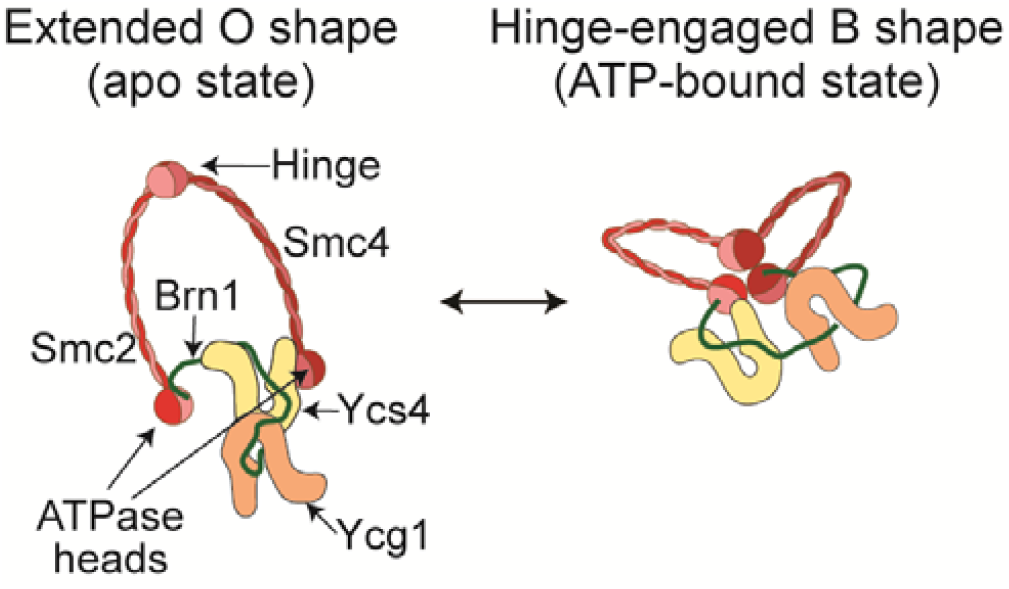
Structural domains of the yeast condensin holocomplex. The condensin holocomplex consists of two antiparallel folded coiled-coil arms, Smc2 and Smc4 (each ~50 nm in length), mutually linked to a hinge domain dimer at one end, and an ATP-binding cassette (ABC)-type nucleotide-binding head domain at the other end, where the two head domains are mutually connected by the Brn1 kleisin subunit. Two HEAT-repeat subunits, Ycs4 and Ycg1, are bound to the Brn1 kleisin subunit. Recent AFM data (18) indicated an O-shape apo state, where the hinge is located far from the globular domain, and an ATP-bound B shape where the hinge is in close proximity of the globular domain (20, 21, 55, 58).

**Supplementary Figure 2.**
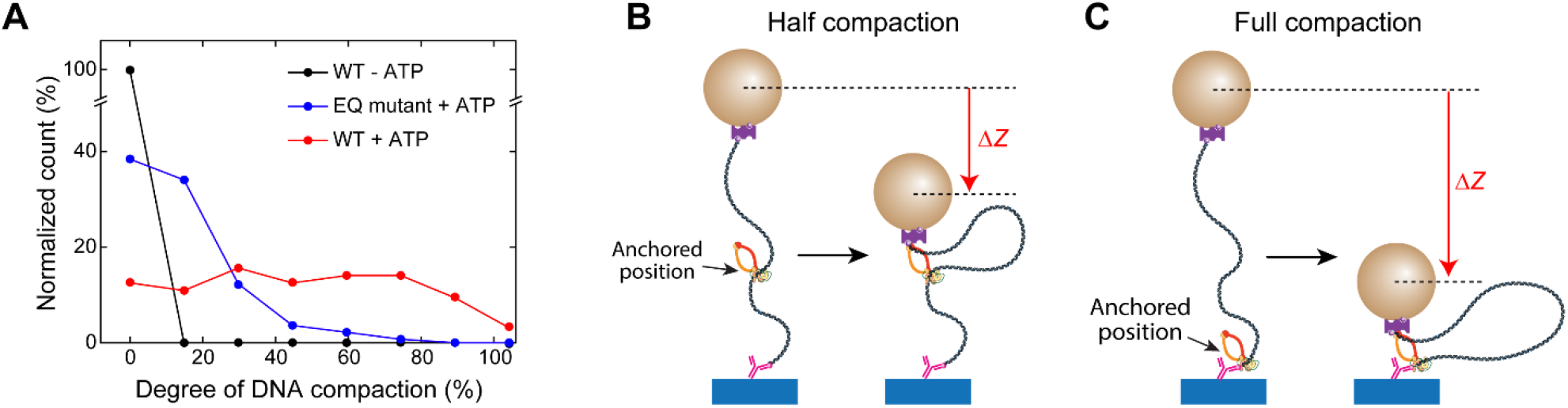
Degree of DNA compaction after DNA loop extrusion activity. **(A)** The degree of DNA compaction (in percent) was measured by the ratio of the bead position immediately after the force jump from 8 pN to 0.2 pN and the bead position after 4 min of DNA loop extrusion (data were taken fromFigure 1B at 0.2 pN (*N* = 58, 91, and 134 in the presence of ATP, in absence of ATP, and for the EQ mutant in presence of ATP, respectively). The average degrees of DNA compaction were 50 ±52 % for the WT in presence of ATP, 0 % for the WT in absence of ATP, and 14 ± 17 % for the EQ mutant in presence of ATP. **(B, C)** Example schematics of DNA compaction when condensin was initially anchored at the middle of the DNA construct (B) and closely at the DNA construct ends (C); these positions lead to half and full DNA compaction, respectively, and illustrate that full compaction is not necessarily expected for the DNA loop extrusion activity of a single condensin complex in this experimental configuration.

**Supplementary Figure 3.**
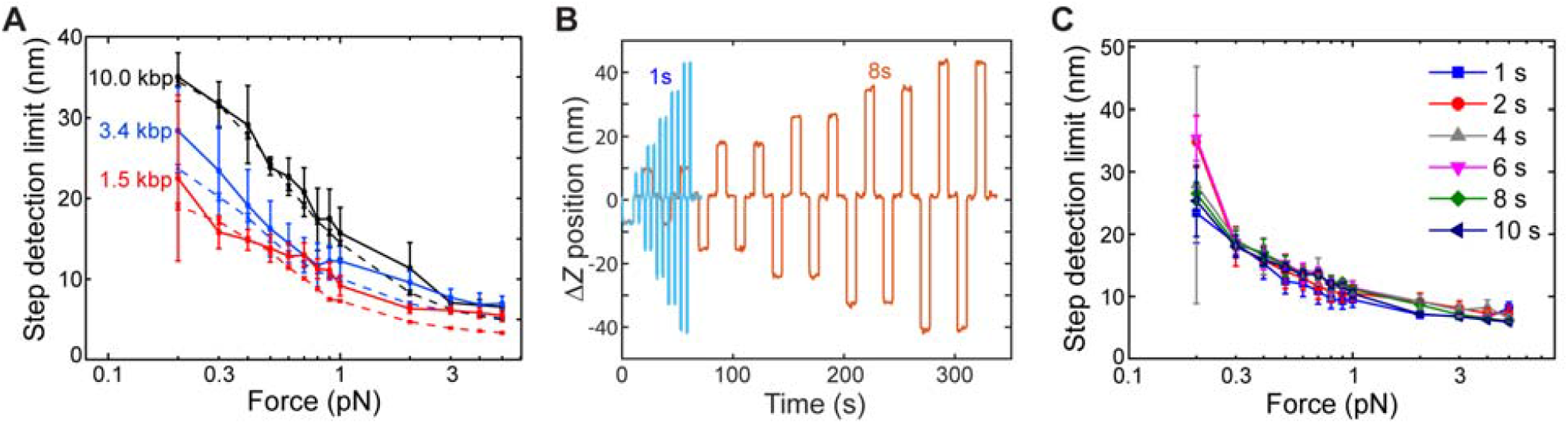
Step-detection limits for different DNA lengths and stretching forces. (**A**) Step detection resolution limit versus DNA stretching force (0.2 – 5 pN) for different DNA lengths (1.5, 3.4, and 10.0 kbp DNA; *N* >80 molecules for each data point) using piezo-induced steps with 10 s plateau (see also Figure 2B). The transverse Δ*Z* bead fluctuations <σ_Z_> (dashed lines) resulting from bead Brownian motion are plotted as well. (**B**) Example trajectories of piezo-induced Δ*Z* steps with 1 s (blue) and 8 s (brown) dwell time between piezo-induced Δ*Z* steps for the 1.5 kbp DNA construct measured at 5 pN. (**C**) Step detection resolution limit versus 1.5 kbp DNA stretching force (0.2 – 5 pN) obtained from different dwell times (1 – 10 s) between the piezo-induced Δ*Z* steps (*N* = 130, 131, 136, 135, 136, 136, 145, 127, 128, 136, 111, 88, 73 for 0.2 pN to 5 pN).

**Supplementary Figure 4.**
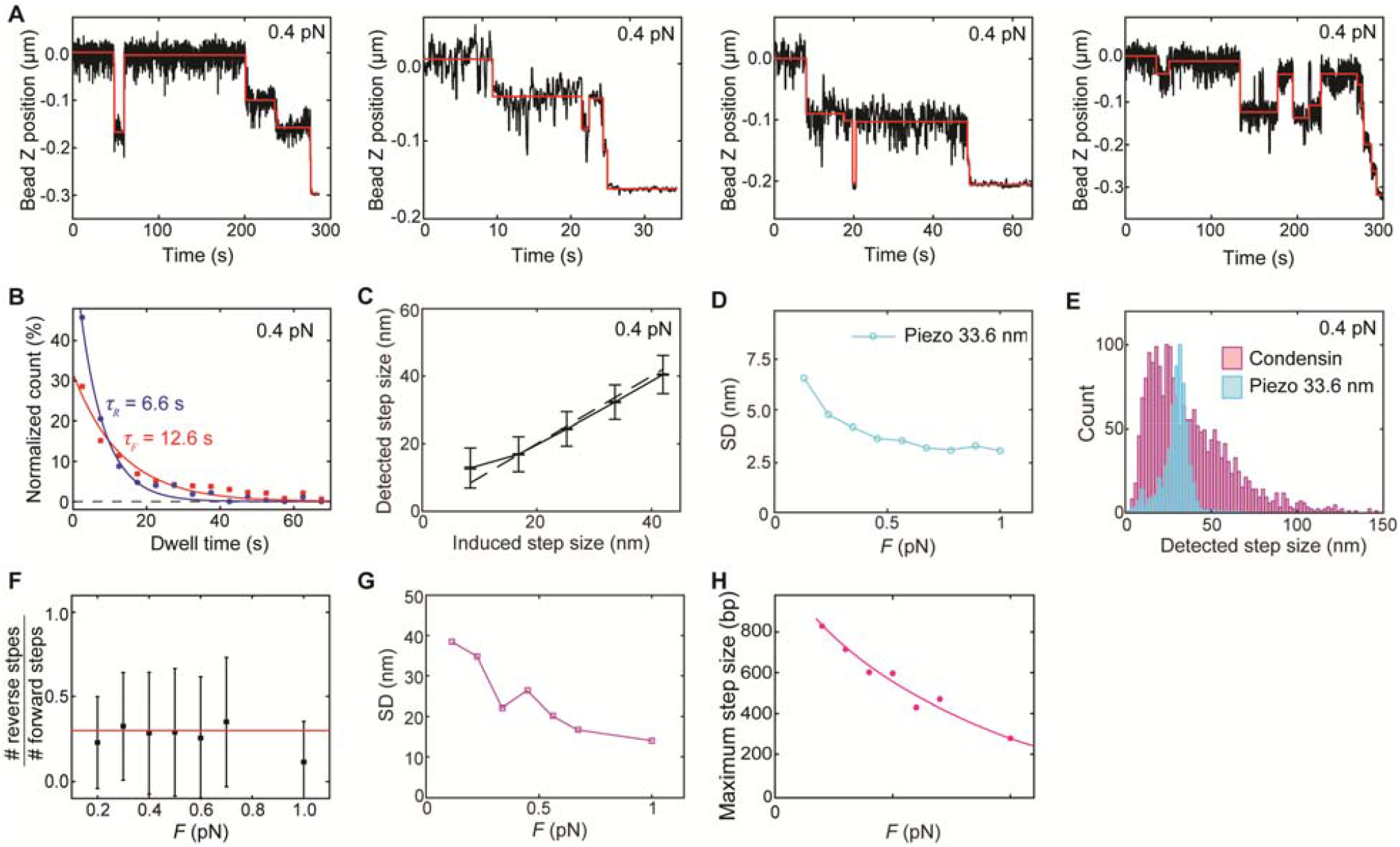
Characterization of steps induced by condensin-mediated DNA loop extrusion and by artificial piezo steps. (**A**) Representative traces of condensin-mediated loop extrusion step sizes at 0.4 pN. Red lines represent fits resulting from the step-finding algorithm. (**B**) Dwell-time distributions of forward (red; *N* = 273) and reverse steps (blue; *N* = 665) that show single-exponential fits (solid lines) with time constants *τ*_*F*_ and *τ*_*R*_, respectively. (**C**) Detected step sizes versus induced step sizes for the different piezo-induced step sizes at 0.4 pN (median ± SD). *N* = 255, 870, 1033, 1069, and 1063 for 8.4, 16.8, 25.2, 33.6, and 42 nm step sizes, respectively. (**D**) Force-dependent width (SD), for the detected step size distributions from piezo-induced artificial step sizes of 33.6 nm (*N* = 994, 1060, 1069, 989, 987, 942, 1,000, 852, and 847 for 0.2, 0.3, 0.4, 0.5, 0.6, 0.7, 0.8, 0.9, and 1 pN, respectively). (**E**) Step size distributions of condensin-mediated DNA loop extrusion (magenta, *N* = 5,128) and piezo-induced artificial step size of 33 nm (cyan, *N* = 1,069) at 0.4 pN. (**F**) The ratio between number of reverse steps and forward steps observed at different DNA stretching forces (mean ± SD; *N* > 500 for each force). Red line depicts the average of the data (y = 0.3). (**G**) Width (SD) of detected step size distributions versus applied force for condensin-induced step (*N* > 500). (**H**) Maximum step sizes obtained from experiments at different forces.

**Supplementary Figure 5.**
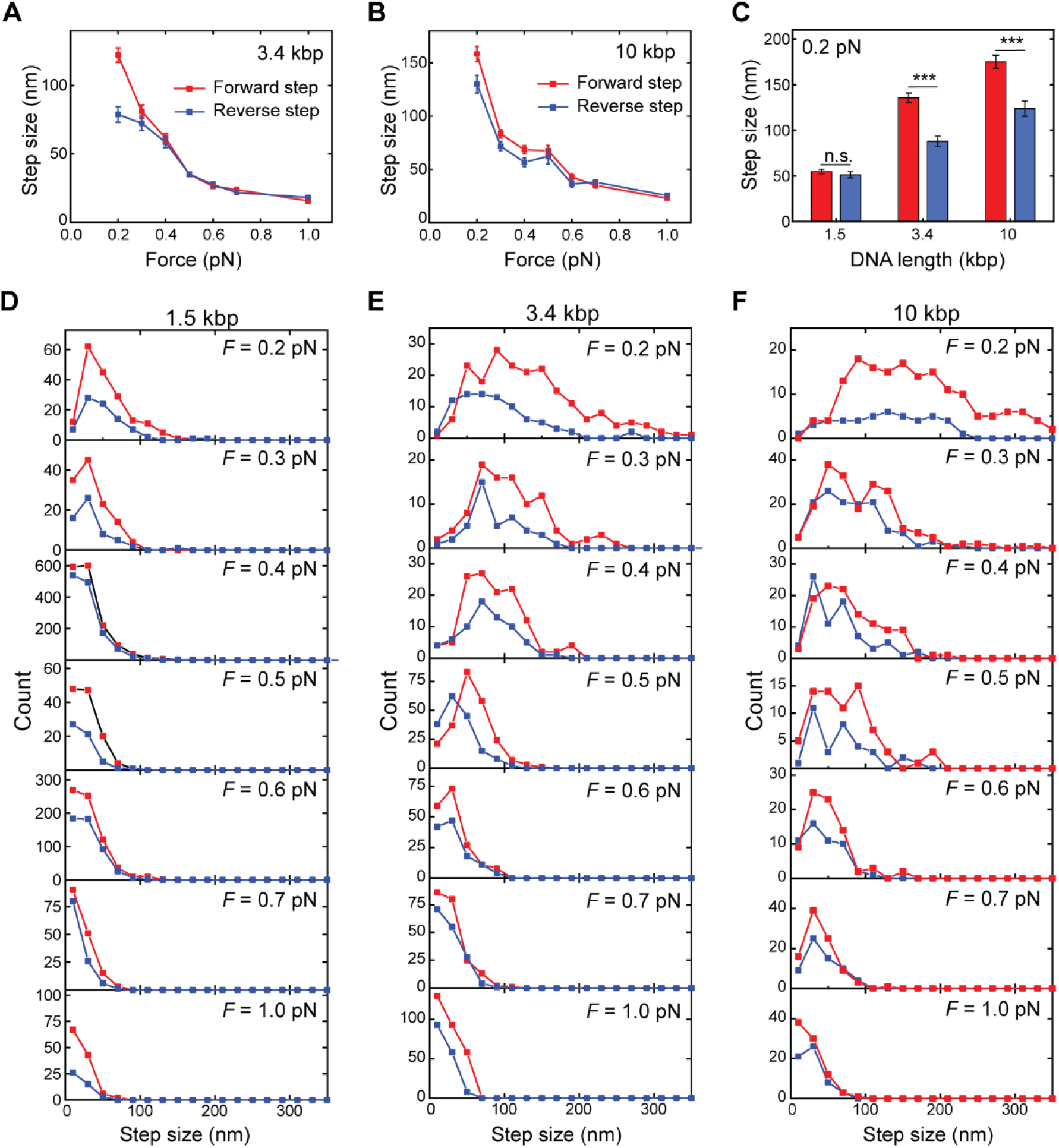
Characterization of forward and reverse steps during DNA loop extrusion for different DNA lengths and at different forces. (**A**) Forward (*N* > 100 for each force) and reverse step (*N* > 40 for each force) sizes in nm (median ± SEM) as a function of force obtained for the 3.4 kbp DNA length. (**B**) Same as (A), but for the 10 kbp DNA length. (**C**) Comparison of average (±SEM) forward (red) and reverse (blue) step sizes at 0.2 pN for the different DNA lengths. Importantly, the forward and reverse step sizes were found to be the same for 1.5 kbp DNA, but significantly different for larger DNA lengths. Statistical analyses consisted of unpaired, two-tailed t-tests (*** indicates p < 0.001; n.s. = non-significant). (**D-F**) Forward (red) and reverse (blue) step size distributions at different forces for DNA lengths of (**D**) 1.5 kbp, (**E**) 3.4 kbp, and (**F**) 10 kbp (*N* > 150 for each histogram).

**Supplementary Figure 6:**
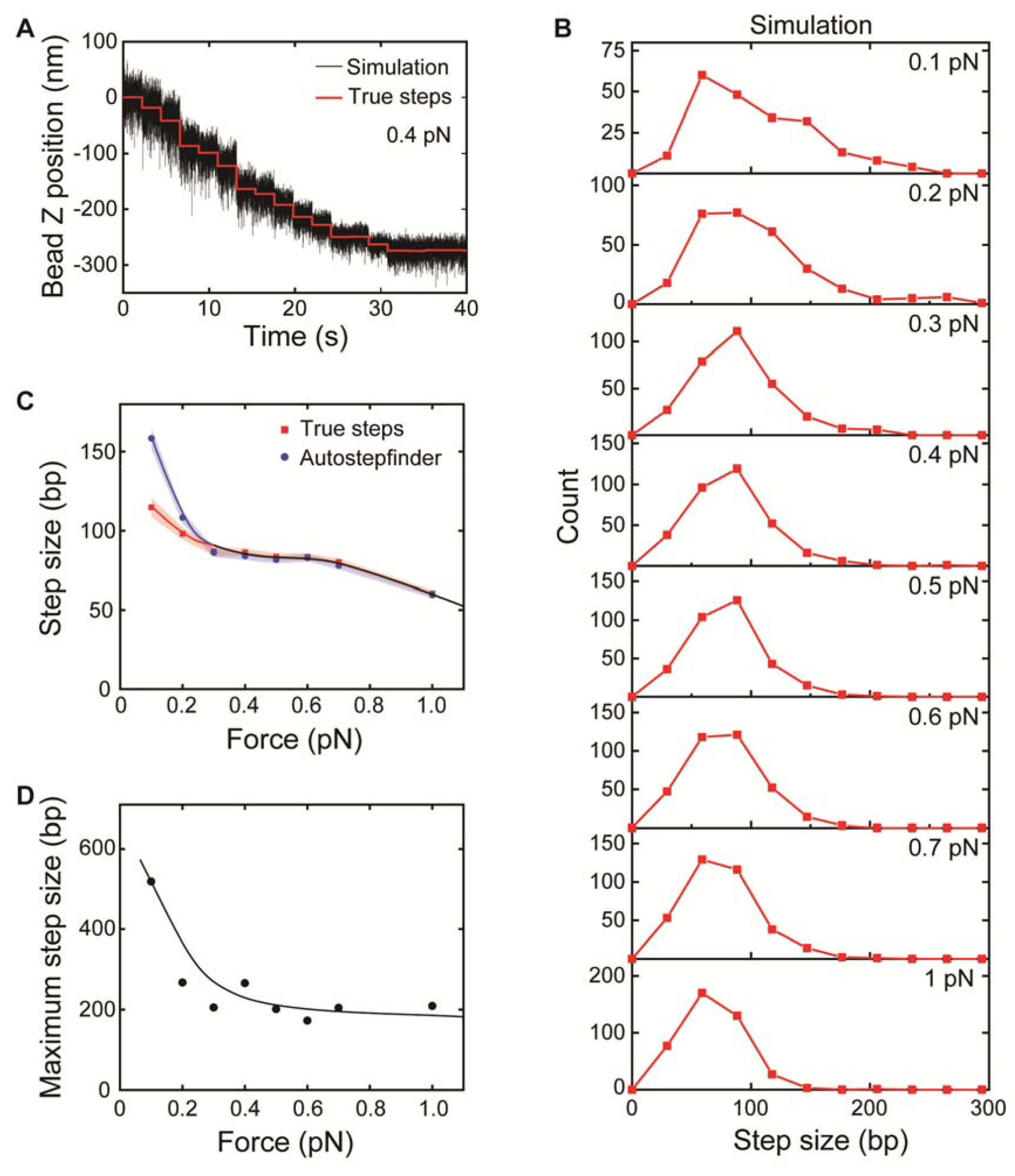
Step size simulation at different forces for the 1.5 kbp DNA construct. (**A**) Representative simulated trace of condensin-mediated DNA loop extrusion at 0.4 pN. Red lines depict true steps (see Material and methods) generated at 2.2 s intervals. (**B**) Simulated step size distributions at different forces (true steps). (**C**) Force-dependent step sizes of simulated step sizes. Step sizes of true steps generated during simulation, and step sizes obtained by the autostepfinder algorithm to simulated traces. (*N* = 213, 291, 305, 329, 328, 355, 353, 409 from 0.1 to 1.0 pN for (B) and (C)). (**D**) Maximum step sizes obtained from simulations at different forces.

**Supplementary Figure 7.**
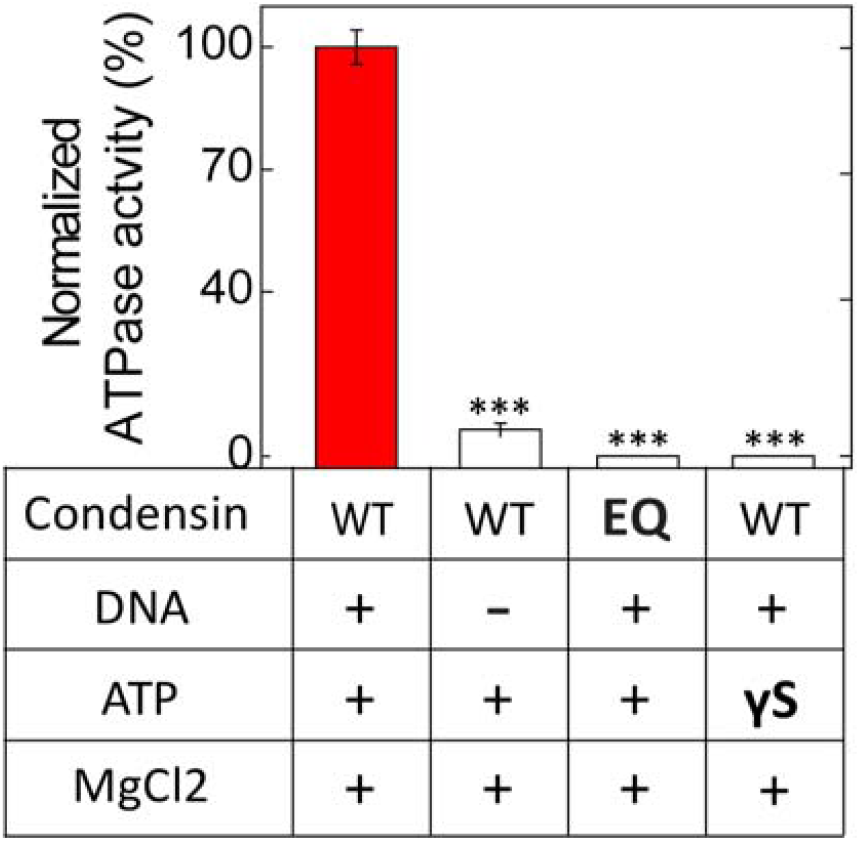
ATPase activity controls for WT condensin and EQ mutant. ATPase activity was measured by the concentration of phosphate ions (*N* = 3 independent experiments). Error bars represent SD. WT denotes wild-type condensin while EQ indicates condensin mutant (Smc2_E1113Q_-Smc4_E1352Q_). γS denotes ATPγS. Statistical analysis consisted of an unpaired two-tailed t-test (*** = *p* < 0.001).

### SUPPLEMENTARY TABLES

**Supplementary Table 1:**
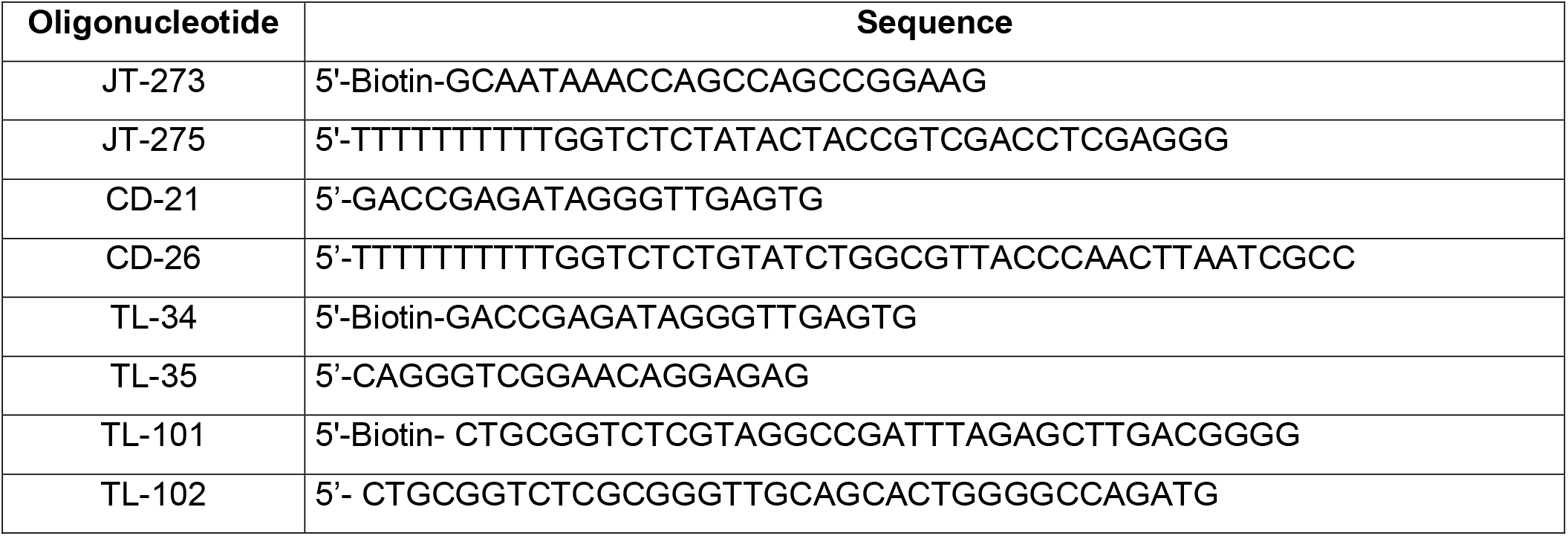
Primer sequences used for the synthesis of linear dsDNA tether constructs.

**Supplementary Table 2:** dsDNA base pair length in nanometer extracted from experimentally determined DNA end-to-end lengths (*N* = 62, 26, and 28 for 1.5, 3.4, and 10 kbp, respectively). Unit is given in nm/bp Persistence length of DNA was 42 nm, resulting from FWLC fit for 1.5 kbp DNA or WLC fit for 3.4 and 10 kbp DNA.

| **Force [pN]** | **1.5 kbp** | **3.4 kbp** | **10 kbp** |
| --- | --- | --- | --- |
| 0.2 | 0.205 | 0.221 | 0.226 |
| 0.3 | 0.231 | 0.243 | 0.246 |
| 0.4 | 0.246 | 0.255 | 0.257 |
| 0.5 | 0.256 | 0.264 | 0.267 |
| 0.6 | 0.264 | 0.269 | 0.271 |
| 0.7 | 0.269 | 0.274 | 0.276 |
| 1 | 0.286 | 0.287 | 0.285 |

